# Exploring Cognitive Individuality and Underlying Creativity in Statistical Learning

**DOI:** 10.1101/2023.07.29.551091

**Authors:** Tatsuya Daikoku, Kevin Kamermans, Maiko Minatoya

**Affiliations:** Graduate School of Information Science and Technology, The University of Tokyo, Tokyo, Japan; Centre for Neuroscience in Education, University of Cambridge, Cambridge, UK; Center for Brain, Mind and KANSEI Sciences Research, Hiroshima University, Hiroshima, Japan

**Keywords:** Phase entrainment, Bayesian, Chunking, Hierarchy, Music, Rhythm

## Abstract

Statistical learning starts at an early age and is intimately linked to brain development and the emergence of individuality. Through such a long period of statistical learning, the brain updates and constructs statistical models, with the model’s individuality changing based on the type and degree of stimulation received. However, the detailed mechanisms underlying this process are unknown.

This paper argues three main points of statistical learning, including 1) cognitive individuality based on “*reliability*” of prediction, 2) the construction of information “*hierarchy*” through chunking, and 3) the acquisition of “1-3Hz *rhythm”* that is essential for early language and music learning. We developed a Hierarchical Bayesian Statistical Learning (HBSL) model that takes into account both reliability and hierarchy, mimicking the statistical learning processes of the brain. Using this model, we conducted a simulation experiment to visualize the temporal dynamics of perception and production processes through statistical learning. By modulating the sensitivity to sound stimuli, we simulated three cognitive models with different reliability on bottom-up sensory stimuli relative to top-down prior prediction: hypo-sensitive, normal-sensitive, and hyper-sensitive models.

We suggested that statistical learning plays a crucial role in the acquisition of 1-3 Hz rhythm. Moreover, a hyper-sensitive model quickly learned the sensory statistics but became fixated on their internal model, making it difficult to generate new information, whereas a hypo-sensitive model has lower learning efficiency but may be more likely to generate new information. Various individual characteristics may not necessarily confer an overall advantage over others, as there may be a trade-off between learning efficiency and the ease of generating new information. This study has the potential to shed light on the heterogeneous nature of statistical learning, as well as the paradoxical phenomenon in which individuals with certain cognitive traits that impede specific types of perceptual abilities exhibit superior performance in creative contexts.

## 1. Introduction

Understanding cognitive individuality and its underlying creativity is crucial for advancing our understanding of human cognition. One critical cognitive function that contributes to language and music acquisition is known as “*statistical learning*” (Saffran et al., 1996). Statistical learning is an innate function of the brain that allows individuals to learn the underlying structure of sensory input by detecting statistical patterns and regularities in the environment.

Recent studies have suggested that individual differences in statistical learning is linked to various cognitive abilities and developmental disorders such as autism spectrum disorder (ASD) and developmental dyslexia (Misyak and Christiansen, 2012; Siegelman et al., 2017; Palmer and Mattys, 2016; Obeid et al., 2016; for review, see Arciuli, 2017; Saffran, 2018). Despite the potential importance of statistical learning in comprehending individual cognitive differences and brain development, it remains unclear how such differences in cognitive abilities arise through statistical learning.

Here, we review neural and computational studies on how cognitive individuality emerges through statistical learning in the brain. Further, for constructive understanding, we conducted a simulation experiment to visualize the temporal dynamics of perception and production processes through statistical learning in different cognitive models. We utilized three models that have varying levels of sensitivity to sound stimuli: hypo-sensitive, normal-sensitive, and hyper-sensitive models. Considering that statistical learning is fundamental to brain development, we also discuss how typical versus atypical brain development influences the perception and production of information through statistical learning.

## 2. Statistical Learning and Its Predictive Processing

### 2.1. Auditory Prediction and Its Individuality

Recently, a growing body of studies has tried to explain the neural and computational mechanisms of learning and generation of auditory structured information (such as music and language) based on the general principle of predictive processing in the brain (Vuust et al., 2022). Predictive processing in the brain works to minimize the prediction error between the bottom-up sensory signals of sound stimuli from the external environment and the top-down predictive signals based on internal models (Friston, 2010; Clark, 2013; Fristone t al., 2017). The “reliability” of the prior probability of top-down predictions is controlled by perceptual uncertainty. The brain learns to adapt continuously to the uncertain environment by reducing perceptual uncertainty and prediction errors.

Researchers have attempted to understand cognitive individuality from the perspective of predictive processing in the brain. For example, it has been explained by the dependence on top-down predictions based on the prior probability of internal models (hypo-/hyper-prior) and the dependence on bottom-up sensory signals from the external environment (hypo-/hyper-sensitive) (Pellicano and Burr, 2012; Philippsen and Nagai, 2022). Intuitively, hyper-prior/hypo-sensitive individuals can be characterized as those with strong judgments based on past experiences and hypo-prior/hyper-sensitive as those who adapt quickly to new environments. Recent studies have suggested that such distinct dependence on prior prediction reflects the dynamics of brain development (Philippsen and Nagai, 2019; Philippsen, Tsuji and Nagai, 2022). Neurotypical children tend to exhibit unstable dependence on prior predictions, but over time, they develop the ability to effectively combine sensory information with prior predictions. This enhances their resilience to disruptions in an uncertain environment. On the other hand, individuals with ASD may exhibit distinct patterns of development in predictive processing (see Table 1). That is, they tend to exhibit stronger dependence or reliance on prior predictions in certain situations (hyper-prior) (Philippsen and Nagai, 2019), while in other circumstances, they may exhibit a weaker dependence or reliance on prior predictions (hypo-prior) or stronger reliance on sensory input (hyper-sensitive) (Sinha et al., 2014; Thye et al., 2018; Robertson et al., 2017). Thus, they tend to exhibit variability in their reliance on prior predictions, rather than a consistent pattern of either enhancement or decrease.

**Table 1.** Individual differences of prediction/sensitivity to external stimuli.

| **Types** | **Individuality** | **References** |
| --- | --- | --- |
| Hypo-prior | Autism spectrum disorder<br>Child | Pellicano et al., 2012<br>Philippsen et al., 2022 |
| Hyper-prior | Child | Philippsen et al., 2022 |
| Hypo-sensitivity | Synaesthesia<br>Attention Deficit<br>Hyperactivity Disorder<br>Affective disorders | Ward et al., 2017<br>Panagiotidi et al., 2018<br>Bijlenga et al., 2017<br>Engel-Yeger et al., 2016<br>Thompson et al., 1990 |
| Hyper-sensitivity | Autism spectrum disorder<br>Synaesthesia<br>Attention Deficit<br>Hyperactivity Disorder<br>Affective disorders<br>Bipolar disorder | Van de Cruys et al., 2019<br>Ward et al., 2017<br>Ward et al., 2017<br>Panagiotidi et al., 2018<br>Bijlenga et al., 2017<br>Engel-Yeger et al., 2016<br>Nathan et al., 1999<br>Lewy et al., 1985<br>Hallam et al., 2009<br>Lam et al., 1990 |

### 2.2. Statistical Learning of Structured Sequence

The statistical learning is an essential cognitive function that is closely linked to brain development (Saffran, 2018) and important for understanding individual differences in the perception and production of music and language within the framework of the predictive processing. The basic mechanism involves calculating the statistical probability of environmental information (particularly the transition probability of sequential information) and the uncertainty of probability distribution, and predicting future information based on an internal probabilistic model acquired through statistical learning. The transition probability is a conditional probability of an event en+1, given the preceding n events based on Bayes’ theorem: *P(e_n+1_|e_n_)*, while the uncertainty is often calculated using several ways of information-theoretical entropy such as conditional entropy: −ΣP(xi)ΣP(xi+1|xi)log2P(xi+1|xi). From the psychological standpoint, the formula can be construed as positing that the brain expects a forthcoming event en+1 based on the most recent preceding events en in a given sequence.

The prediction strategy and resulting impression can vary depending on uncertainty, even when the transition probabilities are identical. For example, a recent neural study has revealed that the brain strategically alters the “*order*” of transition probabilities, that is, the length of the preceding n events used as a reference for expectation, based on the uncertainty of sequential information (Daikoku et al., 2023). Another evidence also showed the preference for music stimuli can be understood as a prediction process. That is, it is represented by precision-weighted inverted U curves of the product of the transition probability and uncertainty (Vuust et al., 2012; 2014; Koelsch et al., 2018; Cheung et al., 2019; Gold et al., 2019). Thus, the prediction strategy (order of transition probability) and individual preference is formed by the integration of uncertainty into probability based on individual’s internal model.

Such uncertainty is not universally inherent in music of language per se. Rather, it is “perceptual” uncertainty that is shaped by an individual’s auditory experience. For example, in the case of language, when a native speaker hears a particular word, the uncertainty in predicting the probable subsequent words is low, making prediction easier. Conversely, for non-native speakers, predicting the next word is more difficult due to higher uncertainty. This is a result of individuals constantly updating their internal models through extended periods of statistical learning, thereby generating an appropriate language probability model. Neural and behavioral studies have highlighted the impact of individual’s auditory experience and expertise on the statistical learning abilities (Daikoku and Yumoto, 2020), which can lead to familiarity with specific types of music or genres (Vuust et al., 2012). Furthermore, in addition to perception and learning, past experiences with statistical learning play a crucial role in the development of individual traits related to music production (composition) and creativity (Daikoku and Yumoto, 2020).

### 2.3. Brain’s Statistical Learning is Bayesian Inference but not Maximum Likelihood Estimation

Importantly, auditory experience affects not only perceptual uncertainty, but also the “reliability” of probabilities. For instance, an A-to-B transition which occurs in (1) 9 out of 10 trials and in (2) 90 out of 100 trials both have a transition probability of 90%. However, the degree of reliability is higher in the former case than the latter. Such reliability is useful for the brain to make judgments even for events with low transition probability. Comparing an event that occurs in 10 out of 100 trials with an event that occurs in 1 out of 10, the brain will recognize that the former is reliably unpredictable and confidently use this information to make predictions.

Neurophysiological studies have observed a gradual representation of statistical learning effects as the number of learning repetitions increases (Daikoku et al., 2015), indicating that the brain’s statistical learning is based on Bayesian inference, which gradually improves the reliability of probabilities with increasing experience rather than on maximum likelihood estimation, which does not vary with learning repetitions. Thus, the amount of learning (auditory experience) not only changes the uncertainty of the brain’s internal model but also the reliability of probabilities, which can affect cognitive individuality and the way of predictions.

However, most studies of statistical learning have referred maximum likelihood estimation based on Markov models or n-gram models that do not consider the “reliability” of probabilities, and thus, have not taken into account the effect of learning trial. Therefore, this study developed a novel model, referred to as a “Hierarchical Bayesian Statistical Learning (HBSL)” model incorporating the Bayesian reliability of probabilities into a Markov model. We then used this model to examine the learning process when a specific auditory stimulus sequence is repetitively learned.

It is of note that the reliability of probabilities is not only subject to the amount of learning (experience), but also to prediction biases. As mentioned above (section 2.1), cognitive individuality can be characterized as those with strong or weak dependence or reliance on the prior probability of internal models, referred to as hyper-prior or hypo-prior, respectively, and those with strong or weak dependence on the auditory inputs, referred to as hyper-sensitive or hyper-sensitive, respectively (Pellicano et al., 2012; Philippsen and Nagai, 2022). Thus, the reliability of prediction can also vary depending on the way of prediction as well as auditory experience. In summary, cognitive individuality is associated with a mixture of “dependency on prior prediction (or sensory signal)” and “amount of statistical learning”.

### 2.4. Hierarchy of Syntactic and Rhythm Structure, and Phase Entrainment

Statistical learning has basically been derived from a hypothesis that explains the mechanism of chunking, which detects information units with high transition probabilities from sequential information such as words or phrases (Saffran et al., 1996). Therefore, many previous studies have examined the neural and computational mechanisms of chunking through statistical learning. On the other hand, recent studies have proposed two types of “*hierarchical*” statistical learning systems (Altman, 2017; Daikoku et al., 2021). The first is a system based on the fundamental function of statistical learning, which groups the chunks of information that have high transition probabilities and integrate them into a cohesive unit. The second is a system that arranges various chunked units to create a hierarchical syntactic structure (Figure 1). That is, statistical learning plays a crucial role in the acquisition of the hierarchy, which is an essential and unique feature of language and music (Patel, 2003).

**Figure 1.**
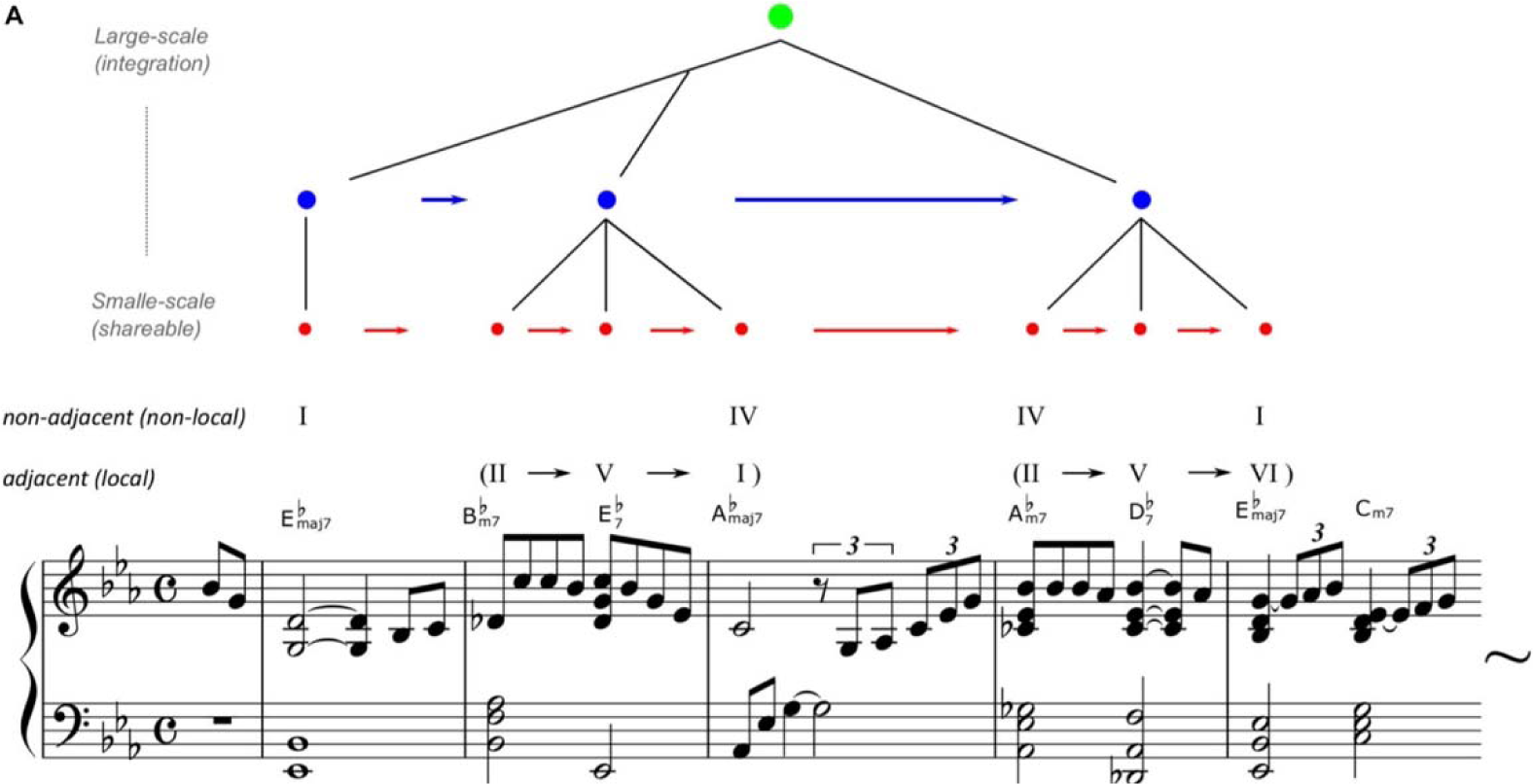
An example of hierarchical statistical learning of music. Reprinted from Daikoku et al. (2021). Misty by Errol Garner, composed in 1954 but arranged by the authors to simplify. The arrangement, chord names, and symbols are simplified (just major/minor, flat, and 7th note) to account for the two-five-one (II–V(7)–I) progression. For example, jazz music has general regularities in chord sequences such as the so-called “two-five-one (II–V– I) progression.” It is a succession of chords whose roots descend in fifths from the supertonic (II) to dominant (V), and finally to the tonic (I). Such syntactic progression frequently occurs in music, and therefore, the statistics of the sequential information have high transitional probability and low uncertainty. Thus, once a person has learned the statistical characteristics, it can be chunked as a commonly used unit among improvisers. In contrast, the ways of combining the chunked units are different between musicians.

Particularly, the hierarchical structure of auditory rhythms has been considered important for the acquisition of music and language (Goswami, 2017). The hierarchy of rhythms refers to a structure in which the lower-level rhythms, such as those corresponding to syllables and musical notes (e.g., crotchet) around 4-12Hz, are included in the higher-level rhythms around 1-3Hz, which correspond to prosody, intonation, and long musical note such as minim (Daikoku et al., 2022). Furthermore, there are rhythms around 12-30Hz that correspond to phonemes or sound onsets at even lower levels of the hierarchy. This can be visualized by analyzing the amplitude modulation (AM) envelope of sound waveforms (Figure 2).

**Figure 2.**
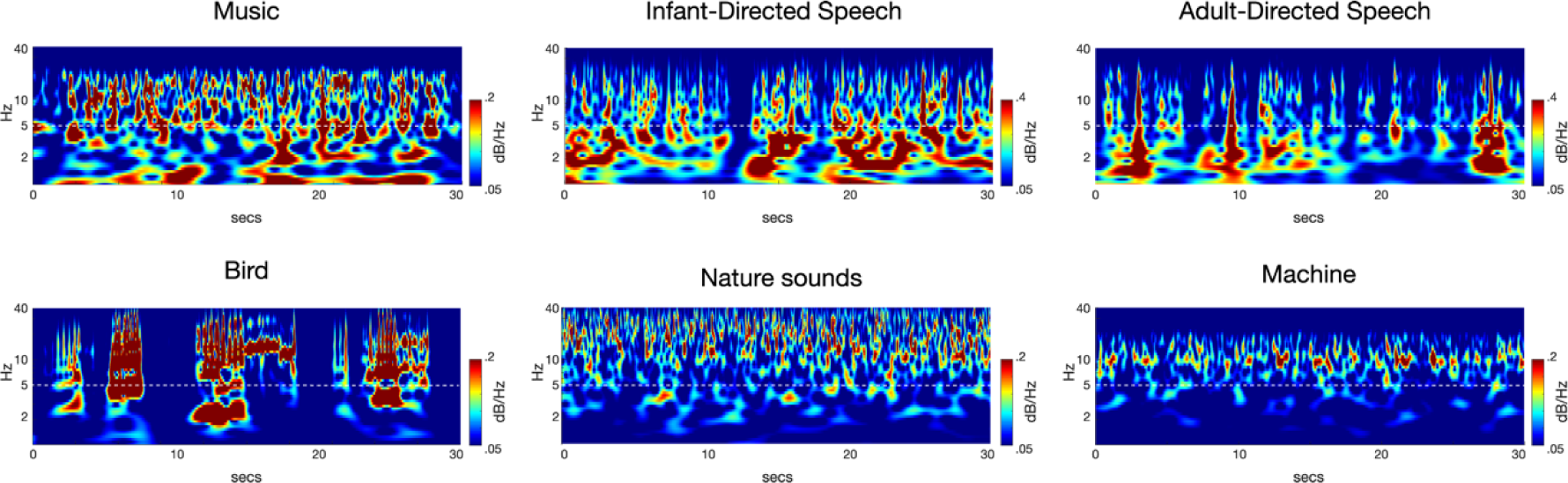
An Example of Rhythm Hierarchy with a Scalogram. Reprinted from the paper by Daikoku and Goswami (2022). Both language and music consist of hierarchical rhythmic structures that include rhythms around 2 Hz, as compared to other auditory stimuli.

It is known that human auditory perception relies in part on phase entrainment of the AM rhythm patterns in sounds at different timescales simultaneously. Such a phase entrainment (also described as phase alignment, neural coupling, tracking, and synchronization) has been shown to contribute to parsing of the sound signal into units such as syllables and words (Poeppel, 2003).

A recent study has shown that the acquisition of the slower rhythm (1-3Hz), that is, phase entrainment of 1-3Hz rhythm (Attaheri et al., 2022) is particularly important for early learning and development of language and music. Notably evidence has also shown that the ability of 1-3 Hz phase entrainment is associated with statistical learning capacity (Assaneo et al., 2019), and neural oscillations synchronize with the statistical chunks acquired via statistical learning (Batterink et al., 2017).

However, brain development can interfere with this function of phase entrainment through statistical learning (Smalle et al., 2022). Individuals with developmental disorders such as Autism Spectrum Disorder (ASD) and developmental dyslexia, which is characterized by difficulties in reading, spelling, and impaired phonological processing (Ramus et al., 2003; Vellutino et al., 2004), exhibit decay of statistical learning and rhythm processing (Arciuli, 2017; Saffran, 2018; Goswami, 2019). Therefore, statistical learning plays a critical role in brain development and the emergence of cognitive individuality. Over a prolonged period of statistical learning, the brain updates and constructs statistical models, with the model’s individuality changing based on the type and degree of stimulation received. However, the detailed mechanisms underlying this process remain unknown.

To provide a constructive understanding of the potential relationships between statistical learning and 1-3Hz rhythm acquisitions, in the next section, we conduct a simulation experiment to visualize the temporal dynamics of perception and production processes through statistical learning, using a newly devised model referred to as the HBSL model with different dependence or reliability on bottom-up sensory stimuli relative to top-down prior prediction: hypo-sensitive, normal-sensitive, and hyper-sensitive models that takes into account both reliability and hierarchy, mimicking the statistical learning processes of the brains with different cognitive individuality. Then, we discuss how atypical cognitive development and individuality (i.e., hypo- and hyper-sensitive) influences the perception and production through statistical learning.

## 3. Simulation

### 3.1. Hierarchical Bayesian Statistical Learning Model

This study developed a computational model, which simulates statistical learning processes of the brain, referred to as HSBL model (Daikoku et al., PAT.P, 2022, Daikoku et al., 2021) (Figure 1). The scripts of the model have been deposited to an external source (https://osf.io/zjwxe/?view_only=4a2f14edd00c4ca391d8befe2e646c73). This is a model that integrates Bayesian estimation with Markov processes using a Dirichlet distribution as a prior distribution. This model can not only calculate the transition probabilities but also determine the “reliability of probabilities” from the inverse of the variance of the prior distribution of the transition probabilities. Using the normalized values of transition probabilities and reliability, this model chunks transition patterns when the product of “reliability * probability” is greater than a constant c. The constant can be decided based on the sample length and the number of learning trial. In this study, we defined c=5 given the sample length used in this experiment. In this study, three models (hypo-, normal-, and hyper-sensitive) with different degrees of dependence on sensory signals were generated by manipulating the parameter vector of the Dirichlet distribution.

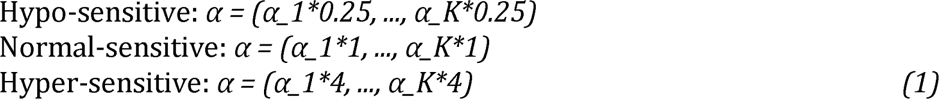

where each α_i corresponds to the prior probability of category i. Specifically, for K categories, α is a K-dimensional vector of positive real numbers. Although the degree of updating transition probabilities remains constant among the three models, differences emerge in terms of the changes in the reliability of transition probability. In the hyper- sensitive model, the reliability of probabilities (variance of prior distribution) varies easily depending on the sensory input, while in the hypo-sensitive model, the reliability of probabilities is less likely to vary even when a new input is provided (eight times weaker than hyper-sensitive model). The normal-sensitive model is an intermediate model between the hyper- and hypo-sensitive models in terms of sensitivity to sensory input.

### 3.2. Learning and Production Processes

We generated fifteen different models by manipulating the degree of dependence on sensory signals and the amount of learning. We used the MIDI data of the Japanese children’s song “Yuuyake Koyake” as the training data, and repeated the learning of the song one to five times using each of the three models (hypo-, normal-, and hyper-sensitive). As a result, a total of fifteen models were generated, consisting of three degrees of dependence on sensory signals and five amounts of learning. We investigate how each of hypo-, normal-, and hyper-sensitive model transforms the internal model over five trials of learning. Furthermore, using the probability distribution of these fifteen models, a hundred pieces of music were probabilistically generated for each model through an automatic composition process (Daikoku et al., PAT.P, 2022).

### 3.3. Comparison of Internal Representations in the Model

We compared the total Bayesian surprise (or total prediction errors) that occurred during learning, measured by the Kullback-Leibler divergence between a distribution P(x) before learning an event (en) and a distribution Q(x) after learning the event (en+1), as well as the total number of chunks generated during 5 trials of statistical learning. The Kullback-Leibler divergence has often been used to measure prediction error or Bayesian surprise in the framework of predictive processing of the brain (Friston, 2010; Baldi and Itti, 2010; Itti and Baldi, 2009). It is a metric used to measure the similarity between two different probability distributions. It represents how much information is lost when one probability distribution changes into another, and since it is non-negative, a small value indicates that the two distributions are similar. Specifically, it is calculated by taking the difference between the probability density functions of the two distributions, taking the logarithm at each point, and then computing the weighted average with respect to one of the distributions. The Kullback-Leibler divergence between two probability distributions P(x) and Q(x) is calculated using the following formula:

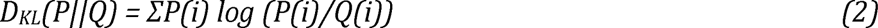

Here, P(i) and Q(i) represent the probabilities of selecting the value i according to the probability distributions P and Q, respectively. In addition, we calculated the average probability distribution of the 100 songs generated by each model and compared the similarity of the models to the training data (i.e., original data) using t-distributed stochastic neighbor embedding (tSNE).

### 3.4. Comparison of Acoustic Properties of Rhythm

We converted the MIDI data of the 100 songs generated by each model into WAV format and extracted the rhythm waveform (modulation wave) below 15 Hz using the Bayesian probabilistic amplitude modulation model (PAD, (Turner and Sahani, 2011)). The acoustic signals were first normalised based on the z-score (mean = 0, SD = 1) in case the sound intensity influenced the spectrotemporal modulation feature. The spectrotemporal modulation of the signals was analysed using PAD to derive the dominant AM patterns. Music and speech signals can be decomposed into slow-varying AM patterns and rapidly-varying carrier or frequency modulation (FM) patterns (Elliott and Theunissen, 2009; Turner, 2010; Daikoku et al., 2022). AM patterns are responsible for fluctuations in sound intensity, which are considered to be a primary acoustic feature of perceived hierarchical rhythm. On the other hand, FM patterns reflect fluctuations in spectral frequency and noise. AM envelopes of speech signals can be separated from the FM structure by means of amplitude demodulation processes. The PAD model employs Bayesian inference to infer the modulators and carrier, and to identify the envelope that best fits the data and a priori assumptions. More specifically, amplitude demodulation is the process by which a signal (yt) is decomposed into a slowly varying modulator (mt) and a rapidly varying carrier (ct):

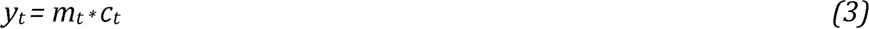

PAD employs amplitude demodulation as a process of both learning and inference. Learning involves the estimation of parameters that describe distributional constraints, such as the expected timescale of variation of the modulator. Inference involves estimating the modulator and carrier from the signals based on learned or manually defined parametric distributional constraints. This information is probabilistically encoded in the likelihood function *P(y_1:T_|c_1:T_, m_1:T_, θ)*, the prior distribution over the carrier *p(*c_1:T_*|θ)*, and the prior distribution over the modulators: *p(m_1:T_|θ)*. Here, the notation x1:T represents all the samples of the signal x, ranging from 1 to a maximum value T. Each of these distributions depends on a set of parameters θ, which control factors such as the typical timescale of variation of the modulator or the frequency content of the carrier. In more detail, the parametrised joint probability of the signal, carrier, and modulator is:

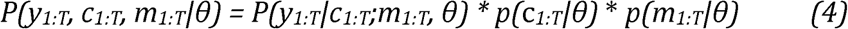

Bayes’ theorem is applied for inference, forming the posterior distribution over the modulators and carriers, given the signal:

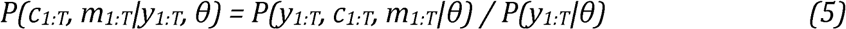

The full solution to PAD is a distribution over the possible pairs of modulators and carriers. The most probable pair of modulator and carrier given the signal is returned:

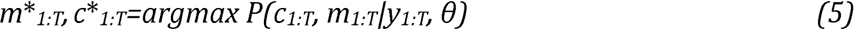

PAD utilizes Bayesian inference to estimate the most suitable modulator (i.e., envelope) and carrier that best align with the data and a priori assumptions. The resulting solution takes the form of a probability distribution, which describes the likelihood of a specific setting of modulator and carrier given the observed signal. Thus, PAD summarizes the posterior distribution by returning the specific envelope and carrier with the highest posterior probability, thereby providing the best fit to the data.

PAD can be run recursively using different demodulation parameters each time, producing a cascade of amplitude modulators at different oscillatory rates to form an AM. The positive slow envelope is modeled by applying an exponential nonlinear function to a stationary Gaussian process, resulting in a positive-valued envelope with a constant mean over time. The degree of correlation between points in the envelope can be constrained by the timescale parameters of variation of the modulator (i.e., envelope), which can either be manually entered or learned from the data.

In the present study, we manually entered the PAD parameters to produce the modulators at an oscillatory band level (i.e., <10 Hz) isolated from a carrier at a higher frequency rate (>10 Hz). The carrier reflects components, including noise and pitches, for which the frequencies are much higher than those of the core modulation bands. In each sample, the modulators (envelopes) were converted into time-frequency domains using scalogram (Figure 2). The scalograms depict the AM envelopes derived by recursive application of probabilistic amplitude demodulation. We then calculated the average frequency power at each frequency and further averaged it over the 100 songs generated by each model.

## 4. Representation of Individual Difference

### 4.1. Learning Process

This study suggests that the Hypo-sensitive model had the highest total Bayesian surprise or total prediction error (i.e., Kullback-Liebler divergence) during learning, followed by Normal-sensitive model and Hyper-sensitive model (Figure 3, left). Furthermore, the Normal-sensitive and Hyper-sensitive models showed a gradual decrease in Bayesian surprise and an increase in chunking through the trial of learning, whereas the Hypo-sensitive model showed no decrease in Bayesian surprise or increase in chunking (Figure 3).

**Figure 3.**
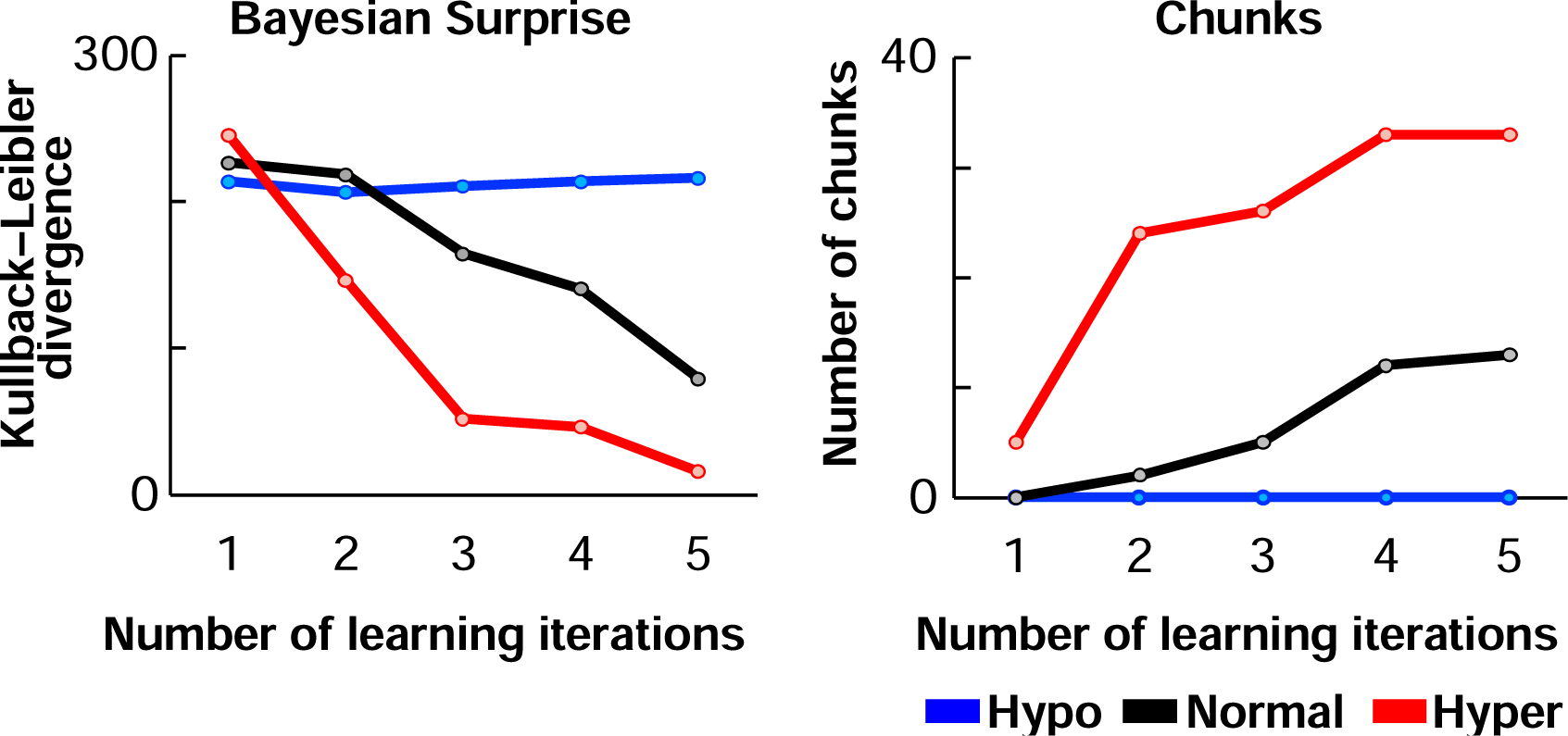
Statistical learning effects in each trial of learning. The left is the total Bayesian surprise (or prediction error) in learning a piece of music, and the right is the number of chunks after statistical learning. Blue, black, and red represent the hypo-, normal-, hyper- sensitive models, respectively.

### 4.2. Production Process

In terms of acoustic features of composed music after learning, both the Hyper- sensitive model and Normal-sensitive model showed a gradual increase in the 2Hz rhythm, which corresponds to short phrases that are considered important in the initial learning of auditory sequences (such as music or language) (Figure 5). On the other hand, rhythms corresponding to notes or beats in the 3-5Hz range gradually decreased with learning. In contrast, the Hypo-sensitive model showed a gradual decrease in the 2Hz rhythm and a gradual increase in the 3-5Hz rhythm with learning.

Regarding probability distribution, the tSNE analysis showed that the similarity of the probability distribution of the composed music to the original music was highest for the Hyper-sensitive model, followed by the Normal-sensitivity model and the Hypo-sensitive model, in that order (Figure 4).

**Figure 4.**
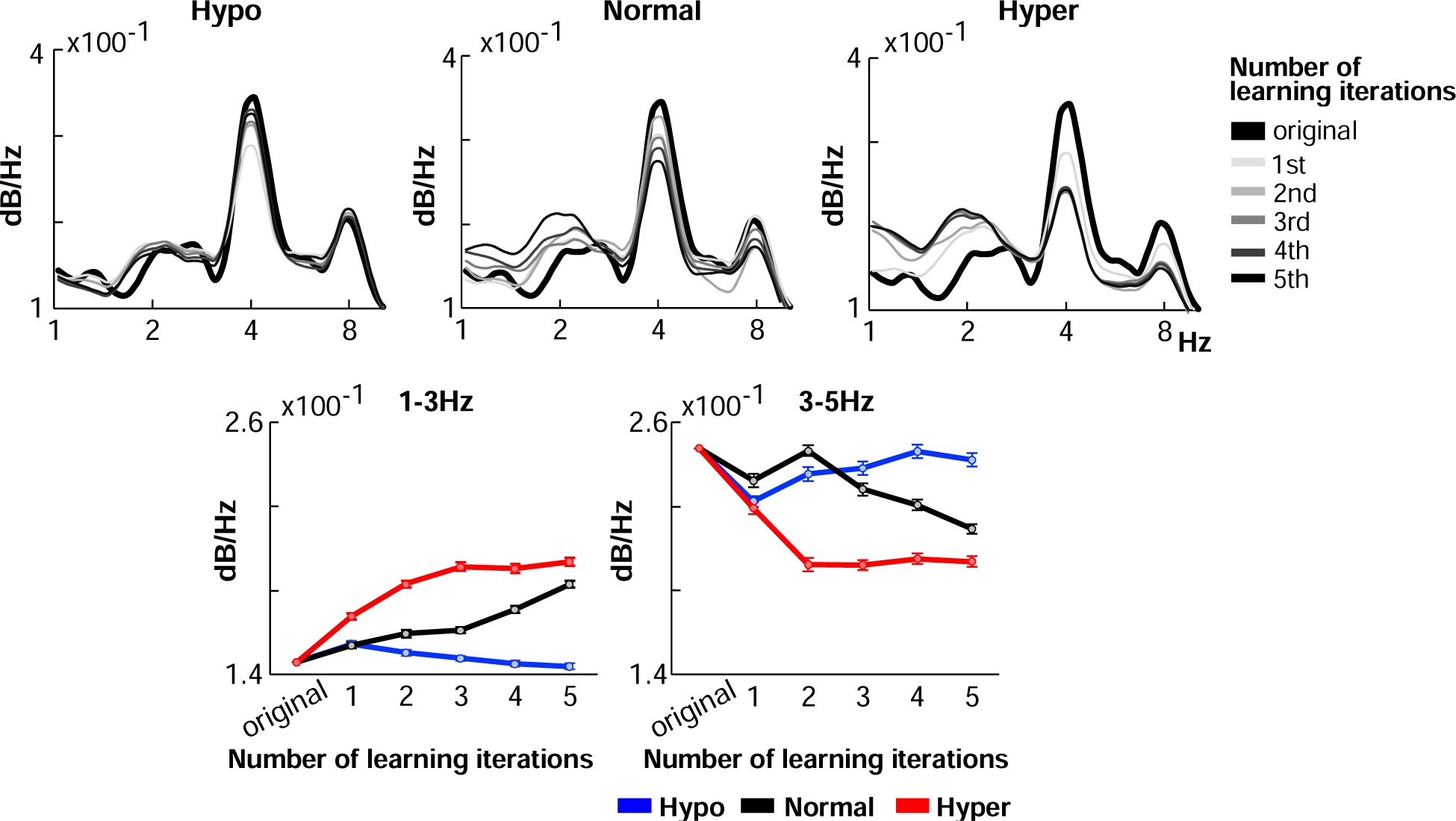
Acoustic properties of composed music after each trial of Statistical learning. Blue, black, and red represent the hypo-, normal-, hyper-sensitive models, respectively.

## 5. Discussion

### 5.1. Emergence of Individuality Through Statistical Learning

Statistical learning is a fundamental process for brain development and contributes to forming individual difference of perception and production (Siegelman et al., 2017; Daikoku et al., 2020; Daikoku, 2018). Computational studies allow for the modeling of the brain’s developmental processes and the emergence of individuality in predictive functions underlying statistical learning. In this study, we used a model that mimics brain’s statistical learning processes including hierarchically structured building to investigate how auditory cognitive individuality arises from statistical learning. Specifically, we conducted a simulation experiment to examine the contributions of two factors to auditory cognitive individuality: 1) sensitivity to sensory signals (hypo-, normal-, hyper-sensitive) and 2) amount of statistical learning (number of learning trials). Our results showed that various auditory cognitive individualities can arise depending on differences in both of sensitivity to sensory signals and amount of statistical learning.

In particular, the normal- and hyper-sensitive models gradually reduced Bayesian surprise, increased the number of chunks (Figure 3), and generated 1-3Hz rhythm (Figure 4) through learning. Moreover, the effects were more pronounced and earlier in the hyper- sensitive model than in the normal-sensitive model. In contrast, in the hypo-sensitive model, neither the reduction of Bayesian surprise nor the chunking occurred through learning. In sum, the simulation experiment showed that the learning efficiency was highest for the hyper model, lowest for the hypo model, and intermediate for the normal model. Due to its high sensitivity to sensory signals, the hyper-sensitive model may exhibit greater adaptability to external input information, potentially resulting in faster learning rates for external sensory statistics.

On the other hand, the hypo-sensitive model produced music with statistically different characteristics from those of the training data (i.e., original music), compared to the other models (Figure 5). That is, the statistical similarity between the generated music and the original music was highest for the hyper-sensitive model and lowest for the hypo- sensitive model. This suggests that the hypo-sensitive model, which showed poor learning efficiency for external information, may have difficulty in statistical learning and chunking, but may be more likely to generate new information. In contrast, the hyper-sensitive model, which can efficiently learn statistical regularities of external information, may have difficulty in generating new information. This suggests that different levels of dependence or reliance on sensory signals lead to differences in the internal model even for the same learning stimuli (Figure 6), and that these differences also influence performance in creative contexts.

**Figure 5.**
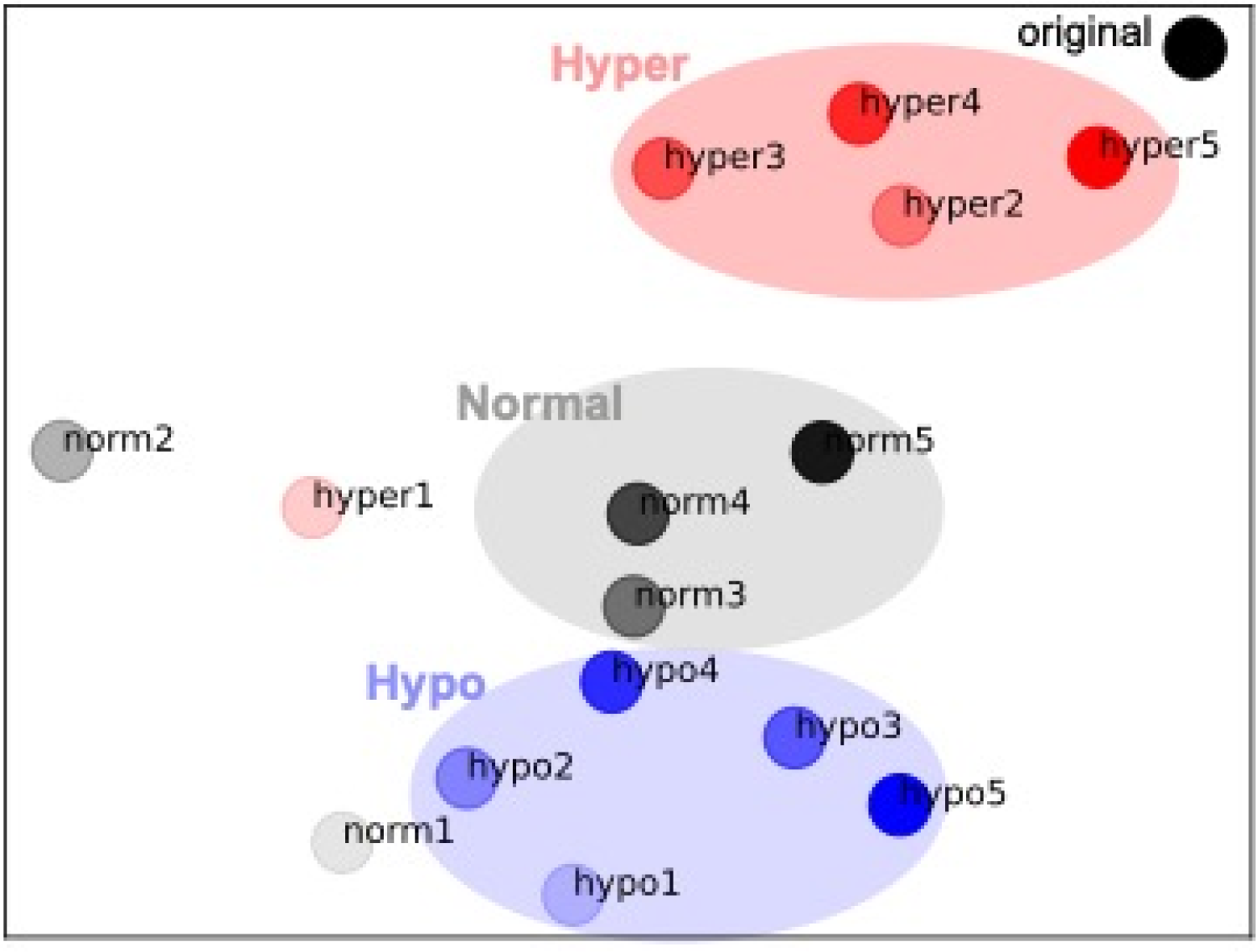
Statistical properties of composed music after each trial of Statistical learning. Blue, black, and red represent the hypo-, normal-, hyper-sensitive models, respectively.

**Figure 6.**
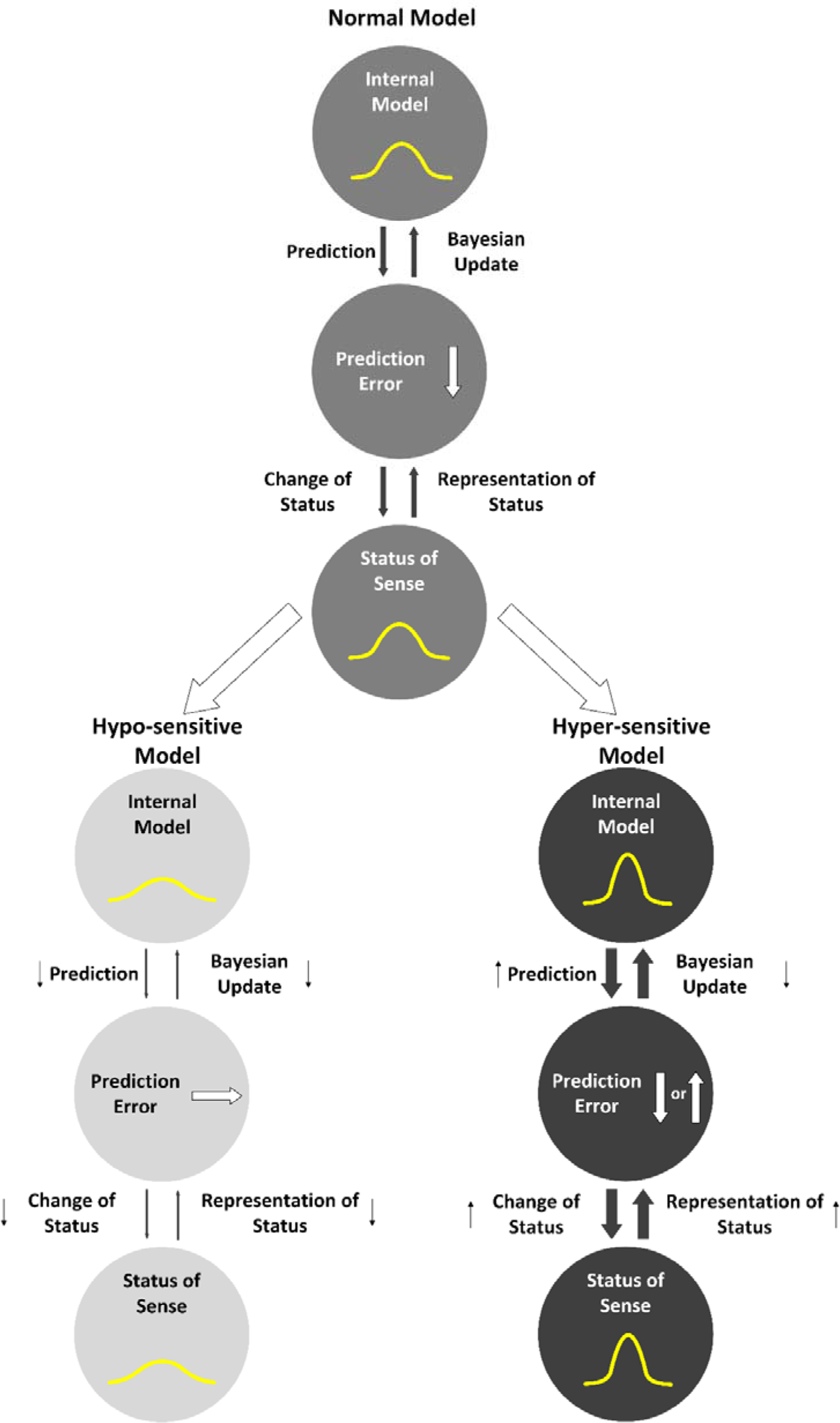
Graphical representation of the normal model, hypo- and hyper-sensitive models

### 5.2. From Statistical Learning to Rhythm Acquisition

This study demonstrated that statistical learning contributes to the acquisition of rhythms around 1-3Hz (Figure 5). Furthermore, while the rhythms around 1-3Hz gradually increased in hyper-sensitive and normal-sensitive models, they gradually decreased in hypo-sensitive. Previous studies have shown that individuals with ASD, who also tend to exhibit hypo-sensitivity, produce speech with weak prosody and monotone intonation, which corresponds to a core 1-3Hz phase component in speech rhythm hierarchy (Kanner, 1943). In addition, a previous study that examined the speech rhythm of individuals with ASD using the PAD model employed in this study revealed a decrease in power around the 1-3Hz frequency range corresponding to the rhythm of prosody and intonation (Daikoku, 2022, PsyArXiv). Taken together with the results of this study, it is possible to say that individuals with hypo-sensitivity may have difficulty acquiring rhythms around 2Hz.

A recent study has shown that the neural processing of the slower rhythm, that is, oscillatory phase entrainment of 1-3Hz rhythm (Attaheri et al., 2022) is particularly important for early learning and development in language. Notably, evidence has also shown that the ability of 1-3 Hz phase entrainment is associated with statistical learning capacity (Assaneo et al., 2019), and neural oscillations synchronize with the statistical chunks acquired via statistical learning (Batterink et al., 2017). However, brain development interfered with this function of phase entrainment in statistical learning (Smalle et al., 2022). Individuals with ASD and developmental dyslexia exhibit decay of statistical learning as well as rhythm processing (Arciuli, 2017; Saffran, 2018; Goswami, 2019). Therefore, statistical learning may play a critical role in acquisition and development of 1-3Hz rhythm.

As stated in the Introduction section, two types of “*hierarchical*” statistical learning systems have been proposed (Altman, 2017; Daikoku et al., 2021). The first is to chunk series of “local” information that have high transition probabilities. The second is to arrange these chunked units to create a “global” hierarchical syntactic structure (Figure 1). In our study, the first function corresponds to the acquisition of 3-5Hz rhythm while the second is the acquisition of 1-3Hz rhythm (Figure 2). According to previous studies, individuals with ASD exhibit inconsistent responses to local deviants (information that induces prediction error). For example, in studies of ASD and mismatch negativity (MMN, an event-related response component in an EEG signal that occurs in response to deviant signals), some studies demonstrated weaker MMN in individuals with ASD than in typically-developed individuals (Seri et al., 1999; Abdeltawwab and Baz, 2014; Bonnet-Brilhault et al., 2016), while other studies detected larger MMN responses in ASD (Gomot et al., 2002, 2011; Ferri et al., 2003; Lepistö et al., 2005; Green et al., 2020). These findings suggest that individuals with ASD may exhibit either hypo-sensitivity or hyper-sensitivity to local sensory properties. However, studies on global predictive processing (e.g., hierarchical structure building) have consistently reported that individuals with ASD exhibit weak response to global deviants (see, Figure 1) (Goris et al., 2018). This implies that ASD is hypo-sensitive to non-local statistics, while sensitivity to local events depends on the type of stimuli (Ide et al., 2017), representing either hypo- or hyper-sensitivity to local statistics.

### 5.3. Balance of Reliance on Prior Prediction and its Development

Past studies suggest that as the brain develops, neurotypical individuals transition from relying heavily on sensory input statistics while giving less weight to prior predictions (known as hypo-prior or hyper-sensitive) to properly integrating sensory statistics with prior predictions (Philippsen and Nagai, 2019). This helps individuals become more resilient in uncertain environments. However, developmental disabilities, such as ASD, may result in different neural mechanisms underlying prior prediction (Nagai, 2019; Lanillos et al., 2020). Studies suggest that individuals with ASD have hyper-plasticity in short-term statistical learning, leading to a preference for recent sensory statistics over global (i.e., long-term) statistical structures (Sinha et al., 2014; Saffran, 2018). This means that individuals with ASD may heavily rely on sensory input while giving less weight to prior prediction (i.e., hypo-prior or hyper-sensitive) during statistical learning. It is worth noting that there may be contrasting abnormalities in predictive function in ASD, with a stronger reliance on prior predictions (i.e., hyper-prior) (Philippsen and Nagai, 2019) instead of hypo-prior predictions (Pellicano and Burr, 2012). Thus, the abnormality of prior prediction in ASD and other developmental disorders may be characterized by instability or variability rather than either enhancement or decay in reliance on prior prediction (for summary, see Table 1).

Several studies have also indicated that children with developmental language disorders, including developmental dyslexia, which is defined by difficulties in reading, spelling, and impaired phonological processing (Snowling, 2000; Ramus et al., 2003; Vellutino et al., 2004), demonstrate a diminished capacity for statistical learning (Daikoku et al., 2023). This suggests that developmental dyslexia may exhibit language impairment resulting from difficulties in detecting and utilizing the statistical regularities of language, leading to a potential hypo-sensitive characterization. Nevertheless, as noted in the features of ASD, they may also display normal predictive processing or hyper-sensitivity to other sensory signals such as music and somatosensory signals. The possibility of varying prediction processing depending on the type of stimulus could be a critical key for future research.

Such instability of reliance on prior prediction could also influence the precision of perceptual uncertainty, as the precision is estimated by the inverse variance of any sensory input (i.e., prior distribution) (Koelsch et al., 2019). Studies have indicated that ASD is susceptible to perceptual uncertainty (Boulter et al., 2014; Lawson et al., 2014; Van de Cruys et al., 2014). The inability to tolerate uncertainty can be considered a key marker of generalized anxiety disorder (Freeston et al., 1994). This trait may also be related to the heightened anxiety commonly observed in people with ASD, and could have a negative effect on their creativity (Baas et al., 2008). It has been suggested that individuals with ASD may experience increased anxiety levels when the level of uncertainty in their environment is high (Boulter et al., 2014). In other words, the fear of uncertain situations can potentially limit the creative potential of individuals with ASD.

However, the unique feature of predictive processing and statistical learning in ASD may not always result in negative outcomes, but could have positive effects in certain situations. Several studies have reported that individuals with ASD sometimes exhibit superiority in certain abilities (Boucher et al., 2012), such as mathematics, visual search skills (O’Riordan et al., 2001), and music and art skills (Happé and Frith, 2009; James, 2010). Therefore, such a unique feature of predictive processing and statistical learning may not always result in negative outcomes, but could have positive effects in certain situations.

Thus, atypical brain development may display specific characteristics (rather than decay or facilitation) of predictive processing. It is assumed that these specificities of predictive processing, that is hypo-/hyper and hypo-/hyper-priors sensitivities, could impact statistical learning ability and (statistical) creativity.

### 5.4. Efficiency in Learning or Novelty in Creation

A previous study has found that individuals with ASD were able to come up with more unconventional and uncertain ideas during divergent thinking tasks compared to typically developed individuals. However, the total number of ideas generated by individuals with ASD was fewer than that of typically developed individuals (Best et al., 2015).

Neural evidence partially supports this finding, and explains it by the hypo- connectivity between the prefrontal cortex and other regions in brains of individuals with ASD (Belmonte et al., 2004; Just et al., 2004; Courchesne and Pierce, 2005; Green et al., 2020; Just et al., 2012). Prior predictions are mainly generated in the frontal regions and transmitted to sensory areas through synaptic connections (Cope et al., 2017; Park et al., 2018). The connectivity between these regions is critical for conveying prior predictions and creating plausible representations of sensory input. However, in individuals with ASD, the altered connectivity between these regions can lead to a modulation of prior predictions, resulting in the production of uncertain information, known as hypo-prior.

Previous studies have shown that neural entrainment induced by statistical learning is enhanced when the prefrontal cortex is temporarily disrupted using repetitive transcranial magnetic stimulation (rTMS). This suggests that the temporary disruption in prefrontal cortex function may have caused a hypo-prior or hyper-sensitive state in the brain, potentially resulting in improved statistical learning ability. Our simulation experiments have also shown that hyper-sensitivity leads to improved statistical learning ability from all aspects of reduction of prediction error, increase of chunk, and 1-3Hz rhythm acquisition, thereby supporting the findings of these previous studies.

However, it is important to note that our simulation only controlled sensitivity (bottom-up processing), not prior (top-down processing), and the models repeatedly learned the same music. Therefore, in the hyper-sensitive model, the reliability of the internal model inevitably increases due to the repeated learning of the same information. This means that hyper-sensitivity “during learning” could lead to a kind of hyper-prior “during production”. Future study needs to investigate how the efficiency in learning (perception) and novelty in creativity (production) are affected when learning various types of information or when controlling for both sensitivity and prior.

In summary, atypical alterations in prior prediction may display specific cognitive individuality involved in perception and production (or learning and creation) through statistical learning. However, such an individuality may not necessarily be favored over the other, as the efficiency of learning and the ease of creating new information may be partially in a trade-off. This study suggests that simulation experiments using statistical learning may lead to a better understanding of the relationship between learning efficiency and creativity in learning systems that exhibit different levels of dependence on sensory signals. Further research on the cognitive individuality may illuminate the potential diversity in human society.

## 6. Summary

This study suggests that hyper-sensitivity allows for efficient statistical learning of information, but makes it difficult to generate new information, while hypo-sensitivity makes it difficult to learn statistically, but may make it easier to generate new information. Different individual characteristics may not necessarily be favored over the other, as the efficiency of learning and the ease of generating new information may be partially in a trade-off. This study has the potential to shed light on the underlying factors contributing to the heterogeneous nature of the supposedly innate ability of statistical learning that all individuals possess, as well as the paradoxical phenomenon in which individuals with certain cognitive traits that impede specific types of perceptual abilities exhibit superior performance in creative contexts.

## Acknowledgements

This research was supported by Japan Society for the Promotion of Science (JSPS) KAKENHI (22KK0157; 21H05063; 22H05210; 22KK0157), and Japan Science and Technology Agency (JST) Moonshot Goal 9 (JPMJMS2296), Japan. The funding sources had no role in the decision to publish or prepare the manuscript.

## Author Contributions

T.D. conceived the method of experiment and data analyses. T.D, analyzed the data and prepared the figures. K.K. and M.M. surveyed previous literature and compiled it into a table. T.D. wrote the original draft of the manuscript. K.K. and M.M. reviewed and edited the manuscript. All authors finalized the manuscript.

## Data Availability

The scripts for the computational model (Hierarchical Bayesian Statistical Learning: HBSL) and analysis and all data including results have been deposited to an external source (https://osf.io/zjwxe/?view_only=4a2f14edd00c4ca391d8befe2e646c73).

